# Structural Basis for Polθ-Helicase DNA Binding and Microhomology-Mediated End-Joining

**DOI:** 10.1101/2024.06.07.597860

**Authors:** Fumiaki Ito, Ziyuan Li, Leonid Minakhin, Htet A. Khant, Richard T. Pomerantz, Xiaojiang S. Chen

**Affiliations:** Molecular and Computational Biology, Department of Biological Sciences and Chemistry, University of Southern California, Los Angeles, California, 90089, USA; Department of Microbiology, Immunology and Molecular Genetics; California NanoSystems Institute, University of California, Los Angeles, CA90095, USA; Department of Biochemistry and Molecular Biology, Sidney Kimmel Cancer Center, Thomas Jefferson University, Philadelphia, PA, 19107, USA; USC Center of Excellence for Nano-Imaging, Viterbi School of Engineering, University of Southern California, Los Angeles, CA, 90089, USA

## Abstract

DNA double-strand breaks (DSBs) present a critical threat to genomic integrity, often precipitating genomic instability and oncogenesis. Repair of DSBs predominantly occurs through homologous recombination (HR) and non-homologous end joining (NHEJ). In HR-deficient cells, DNA polymerase theta (Polθ) becomes critical for DSB repair via microhomology-mediated end joining (MMEJ), also termed theta-mediated end joining (TMEJ). Thus, Polθ is synthetically lethal with BRCA1/2 and other HR factors, underscoring its potential as a therapeutic target in HR-deficient cancers. However, the molecular mechanisms governing Polθ-mediated MMEJ remain poorly understood. Here we present a series of cryo-electron microscopy structures of the Polθ helicase domain (Polθ-hel) in complex with DNA containing 3′-overhang. The structures reveal the sequential conformations adopted by Polθ-hel during the critical phases of DNA binding, microhomology searching, and microhomology annealing. The stepwise conformational changes within the Polθ-hel subdomains and its functional dimeric state are pivotal for aligning the 3′-overhangs, facilitating the microhomology search and subsequent annealing necessary for DSB repair via MMEJ. Our findings illustrate the essential molecular switches within Polθ-hel that orchestrate the MMEJ process in DSB repair, laying the groundwork for the development of targeted therapies against the Polθ-hel.

## Introduction

The integrity of the genome is central to cellular health and proper organismal development. Among the various types of genomic insults, DNA double-strand breaks (DSBs) are particularly catastrophic, potentially leading to severe genomic disorders and tumorigenesis. Cells predominantly repair these DSB lesions through the major DSB repair mechanisms such as homologous recombination (HR) and non-homologous end joining (NHEJ). HR preserves genomic stability by accurately rejoining broken DNA ends. Thus, major HR factors such as BRCA1/2 act as tumor suppressor proteins^1-3^. However, in cells deficient in HR, typically due to mutations in BRCA1/2, alternative pathways like microhomology-mediated end joining (MMEJ) become crucial. DNA polymerase theta (Polθ) promotes DSB repair via MMEJ, also referred to as theta-mediated end-joining (TMEJ) or alternative end-joining (Alt-EJ)^4-7^. Polθ is upregulated in several cancers and plays a pivotal role in DSB repair in HR-deficient (HRD) cancer cells^6,8-10^, making it a promising target for therapeutic intervention^11-16^.

The Polθ protein, consisting of 2,575 amino acids, includes a helicase domain (Polθ-hel), a central flexible domain (Polθ-Ct), and a polymerase domain (Polθ-pol)^5,17-19^. The Polθ-hel has been shown to promote MMEJ in cells and in vitro, but its mechanistic involvement remains unclear^4,10,20^. Polθ-hel is capable of unwinding DNA in a 3′-to-5′ direction and translocating ssDNA in an ATP-dependent manner^19,21,22^. Polθ-hel’s ATPase activity is most strongly stimulated by ssDNA, and it has been shown to promote ssDNA annealing in an ATP-independent manner. Thus, putative functions for Polθ-hel in MMEJ are translocation along ssDNA and annealing of microhomologies crucial for the error-prone MMEJ repair^18,23-25^. The involvement of Polθ-hel in MMEJ make it a promising target for therapeutic interventions in HRD cancer such as subsets of breast, ovarian, prostate, and pancreatic cancers, and the first Polθ-hel inhibitor has entered clinical trials^4,20,26,27^.

Despite its critical roles in MMEJ and promise as a drug target, the molecular details of how Polθ-hel binds to 3′-ssDNA and facilitates the searching and annealing of DNA microhomologies during MMEJ have not been elucidated. Our work offers new structural and biochemical insights that elucidate detailed mechanisms of Polθ-hel DNA binding and microhomology search and annealing, expanding our understanding of this enzyme’s role in initiating MMEJ. Our findings underscore the intricate mechanisms of Polθ-hel’s role in Polθ-mediated DNA repair, providing a basis for novel therapeutic strategies in HRD malignancies.

## Results

### Architecture of Polθ-hel bound to DNA in multiple states

To understand the mechanisms of DNA binding and initial steps of microhomology search and annealing by Polθ, we reconstituted the helicase domain (Polθ-hel) with model MMEJ substrates containing various sequences at the 3′-ssDNA overhang strand (Fig. 1a-b). Fluorescence anisotropy and gel shift assays showed the DNA substrates readily bind to Polθ-hel with similar affinity (Fig. 1c-d). The reconstituted complexes were subsequently applied to structural analysis using single particle cryoEM. To simulate the forms of DNA being repaired through the cellular MMEJ process, we used DNA oligonucleotides consisting of a 30-base pair duplex and a poly(T) 3′-overhang ssDNA. The 3′-overhang region was designed to have various lengths with or without 6-nucleotide long palindromic microhomology (MH) sequence (CCCGGG) that can be annealed to each other at the end of 3′-overhang strand as previously report^10^ (Fig. 1b). Polθ-hel in complex with DNA encompassing 3′-overhangs of 9-nt poly(T) with MH, 11-nt poly(T) with MH, and 15-nt poly(T) without MH, yielded cryoEM reconstructions with resolutions ranging from 3.1 to 3.8 Å (Table 1, Extended Data Fig. 1-3). In addition to the DNA-bound forms, we obtained structures of Polθ-hel bound to AMP-PNP, a non-hydrolyzable ATP analog, and the apo form of the protein (Table 1, Extended Data Fig. 4-5). Among the helicase domain of Polθ that are included in our construct (amino acid residues 1-894), all the cryoEM reconstructions showed density for the core subdomains comprising the helicase ring, including the two RecA-like domains (D1 and D2), winged-helix (WH) domain (D3), and ratchet domain (D4) (Fig. 1a, e-g). A small helix-loop-helix (HLH) domain (D5) appeared conformationally variable and visible for a subset of the structures (discussed later).

**Fig. 1.**
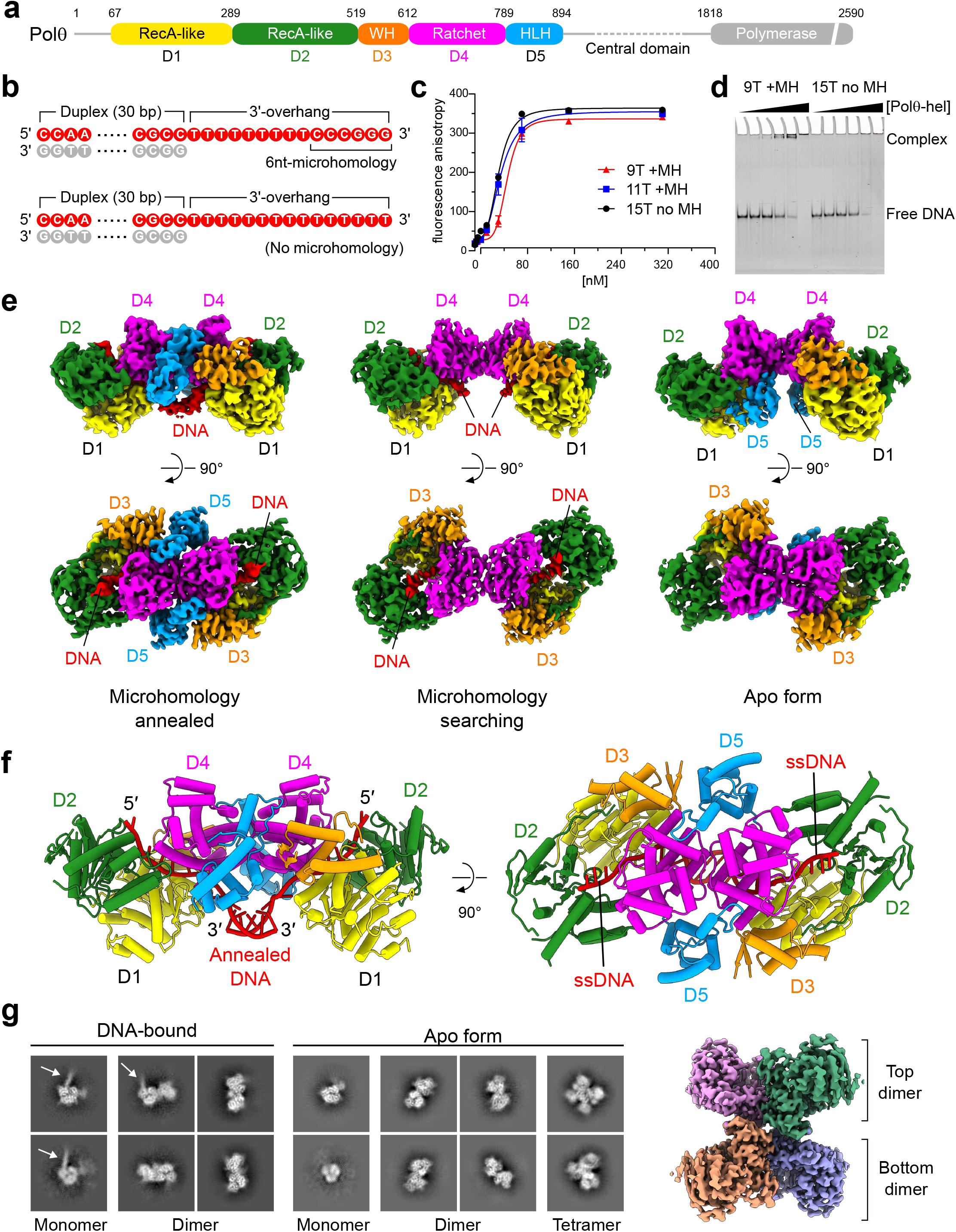
Architecture of Polθ-hel-DNA complex. (**a**) Schematic representation of the domain organization and construct design of Polθ-hel. WH, winged helix; HLH, helix-loop-helix. The regions either invisible in any of the cryoEM maps or excluded from the construct are colored in grey. (**b**) DNA substrate sequences used in this study. A 30-base pair DNA duplex with a poly(T) 3′-overhang ssDNA was prepared either with 6-nt MH sequence (CCCGGG) or without MH sequence. (**c**) Quantitation of DNA binding of Polθ-hel by rotational anisotropy. The FAM-labeled DNA with 3′-overhang ssDNA containing 9-nt poly(T) with MH, 11-nt poly(T) with MH, and 15-nt poly(T) without MH were used for the binding assay. (**d**) DNA gel shift assay with unlabeled DNA with 3′-overhang ssDNA containing 9-nt poly(T) with MH and 15-nt poly(T) without MH. (**e**) Orthogonal views of the cryoEM maps of Polθ-hel dimer in complex with 3′-overhangs with 9-nt poly(T) with MH (left), 15-nt poly(T) without MH (middle), and in apo form (right). (**f**) Orthogonal views of the atomic model of the Polθ-hel in MH annealed state. (**g**) Representative 2D class averages from the data sets of the complex with 15-nt poly(T) DNA without MH 3′-overhang DNA (left) and apo form (middle). The cryoEM map of the apo form tetramer is shown on the right.

**Table 1.** Cryo-EM data collection, refinement, and validation statistics.

|  | Polθ-hel-DNA<br>MH annealed<br>state 1<br>dimer<br>(EMDB: EMD-45217) | Polθ-hel-DNA<br>MH annealed<br>state 2<br>dimer<br>(PDB: 9C5Q)<br>(EMDB: EMD-45218) | Polθ-hel-DNA<br>MH search<br>dimer<br>(PDB: 8W0A)<br>(EMDB: EMD-43706) | Polθ-hel-AMP-PNP<br>dimer<br>(PDB: 9ASJ)<br>(EMDB: EMD-43816) | Polθ-hel<br>apo form<br>dimer<br>(PDB: 9ASK)<br>(EMDB: EMD-43817) | Polθ-hel<br>apo form<br>tetramer<br>(PDB: 9ASL)<br>(EMDB: EMD-43818) |
| --- | --- | --- | --- | --- | --- | --- |
| **Data collection** |  |  |  |  |  |  |
| Magnification | 105,000 | 105,000 | 105,000 | 150,000 | 150,000 | 150,000 |
| Voltage (kV) | 300 | 300 | 300 | 200 | 200 | 200 |
| Electron exposure (e<sup>-</sup>/Å<sup>2</sup>) | 65 | 65 | 65 | 58 | 58 | 58 |
| Defocus range (μm) | -0.6 to -2.0 | -0.6 to -2.0 | -0.6 to -2.0 | -1.0 to -3.0 | -1.0 to -3.0 | -1.0 to -3.0 |
| Pixel size (Å) | 0.86 | 0.86 | 0.86 | 0.92 | 0.92 | 0.92 |
| Symmetry imposed | C1 | C2 | C2 | C2 | C2 | D2 |
| Initial particle images | 885,890 | 3,203,827 | 2,654,648 | 509,731 | 3,961,362 | 3,961,362 |
| Final particle images | 53,607 | 78,928 | 99,902 | 68,248 | 280,269 | 202,004 |
| Map resolution (Å) | 3.75 | 3.13 | 3.23 | 3.54 | 3.61 | 3.51 |
| FSC threshold | 0.143 | 0.143 | 0.143 | 0.143 | 0.143 | 0.143 |
| Map resolution range (Å) | 2.8 – 8.3 | 2.5 – 6.0 | 2.6 – 3.7 | 3.0 – 4.0 | 3.0 – 4.2 | 3.0 – 4.0 |
| **Refinement** |  |  |  |  |  |  |
| Initial model used (PDB) |  | Polθ-hel-DNA<br>MH search | 5AGA | Polθ-hel apo<br>form dimer | 5AGA | Polθ-hel apo<br>form dimer |
| Model resolution (Å) |  | 3.5 | 3.6 | 3.9 | 4.0 | 3.8 |
| FSC threshold |  | 0.5 | 0.5 | 0.5 | 0.5 | 0.5 |
| Map sharpening B factor (Å<sup>2</sup>) |  | -44.0 | -82.5 | -110.1 | -154.6 | -145.4 |
| No. non-hydrogen atoms |  | 12418 | 10216 | 11626 | 11564 | 23128 |
| Protein residues |  | 1550 | 1272 | 1482 | 1482 | 2964 |
| Nucleotides |  | 28 | 14 | 0 | 0 | 0 |
| Ligands |  | 0 | 0 | 2 | 0 | 0 |
| *B*-factors |  |  |  |  |  |  |
| Protein |  | 105.55 | 48.54 | 61.54 | 52.01 | 68.12 |
| Nucleotide |  | 111.01 | 10.40 | - | - | - |
| Ligand |  | - | - | 94.67 | - | - |
| R.m.s. deviations |  |  |  |  |  |  |
| Bond lengths (Å) |  | 0.002 | 0.004 | 0.004 | 0.003 | 0.004 |
| Bond angles (°) |  | 0.555 | 0.972 | 0.969 | 0.604 | 0.681 |
| Validation |  |  |  |  |  |  |
| MolProbity score |  | 1.90 | 2.02 | 1.95 | 1.96 | 2.07 |
| Clash score |  | 7.17 | 10.31 | 13.06 | 14.03 | 16.24 |
| Poor rotamers (%) |  | 0.08 | 0.09 | 0.24 | 0.32 | 0.44 |
| Ramachandran plot |  |  |  |  |  |  |
| Favored (%) |  | 91.50 | 91.94 | 95.37 | 95.64 | 94.81 |
| Allowed (%) |  | 8.50 | 8.06 | 4.63 | 4.36 | 5.19 |
| Disallowed (%) |  | 0.00 | 0.00 | 0.00 | 0.00 | 0.00 |

All the Polθ-hel structures had dimers as the most abundant oligomeric state. The dimerization is mediated by the D4-D4 head-to-head interaction, similar to the previously reported crystal structure of the Polθ-hel tetramer form that contained two copies of this dimer form^24^. In all the DNA-bound structures, clear ssDNA densities were observed along the central channels of the two helicase protomers. The two DNA strands enter through the helicase channel openings that are distantly positioned yet originate from the same side where the subdomain D2 is largely exposed. On the other hand, the 3′-ends of the strands exit the channel on the opposite side of the dimer, bringing the ssDNA ends close to each other (Fig. 1e). In the complex with 3′-overhang DNA with 9-nt poly(T) and 6 nt MH sequence, a continuous low-resolution density spans the dimer cleft and connects the two channel exits for the 3′-end ssDNA (Fig. 1e, top left). The corresponding density in the structure containing poly(T) 3′-overhang without MH was noticeably weaker at a similar isosurface threshold level. Therefore, this density accounts for the microhomology DNA introduced at the ends of the 3′-overhangs. MH annealed DNA was built into the density and termed MH annealed state hereafter (Fig. 1e, left, and 1f). Similarly, the complex with 3′-overhang DNA with 11-nt poly(T) with MH also showed similar densities accounting for the annealed MH at the dimer cleft (Extended Data Fig. 2). In contrast, 3′-overhang DNA with 15-nt poly(T) without MH had a density at the same location, which diminished immediately after exiting the helicase channels, attributable to the absence of MH, with the two 3′-overhangs continuing their search for microhomology, termed the MH search state henceforth.

In addition to the dimer form, the data sets for the MH annealed state and apo form contained tetramer form as a subpopulation, closely resembling the previously reported crystal structure of the tetramer^24^ (Extended Data Fig. 1 and 4). Careful inspection of the cryoEM map of the tetramer form from the sample containing MH DNA showed no trace of DNA density, and it is identical to the tetramer form from the apo form data set. These observations indicate that the tetramer form is incompatible with the DNAbinding. Supporting this observation, the data set for the MH search state structure, where we added an excess amount of DNA (30-fold more than the protein in molarity), yielded 2D class averages predominantly showing dimers and dissociated monomers with no obvious tetramer form. This marks a clear shift from the apo form data set, which exhibited a mixture of tetramer, dimer, and monomer forms (Fig. 1g). Notably, a subset of 2D class averages of the Polθ-hel-DNA showed additional thread-like densities stemming from the Polθ-hel ring, representing the bound DNA (indicated by white arrows in Fig. 1g). These results indicate that the oligomeric state of the Polθ-hel equilibrates between dimer and tetramer, with DNA binding promoting the dissociation of the tetramers into two dimers.

### Polθ-hel D5 is a mobile domain

Unexpectedly, the subdomain D5, composed of short alpha-helices and connecting loops, shows considerable structural variability among our Polθ-hel structures. In the apo form and the AMP-PNP-bound form, D5 adheres to the surface of the helicase ring across the D1-D4 subdomain interface, consistent with the reported crystal structure^24^ (Fig. 2a, top left and middle). Of note, amino acids 839-858 in D5 are disordered in these forms. In the MH search state, the D5 became completely invisible, even at a low isosurface threshold, indicating detachment from the helicase ring and flexible connection via the D4-D5 linker (Fig. 2a, top right). In the MH annealed structures, the D5 reemerges at a different location with a different structure: it is located at a narrow cleft near the D4-D4 dimer interface, surrounded by the D4 of the same protomer and the D3 and D4 of the other protomer (Fig. 2a, bottom). This relocalization results in a large displacement with the C-terminal helix moving approximately 42 Å from its original position in the apo form, accompanied by extensive rotational movements that orient the C-terminal helix in nearly opposite direction (Fig. 2b).

**Fig. 2.**
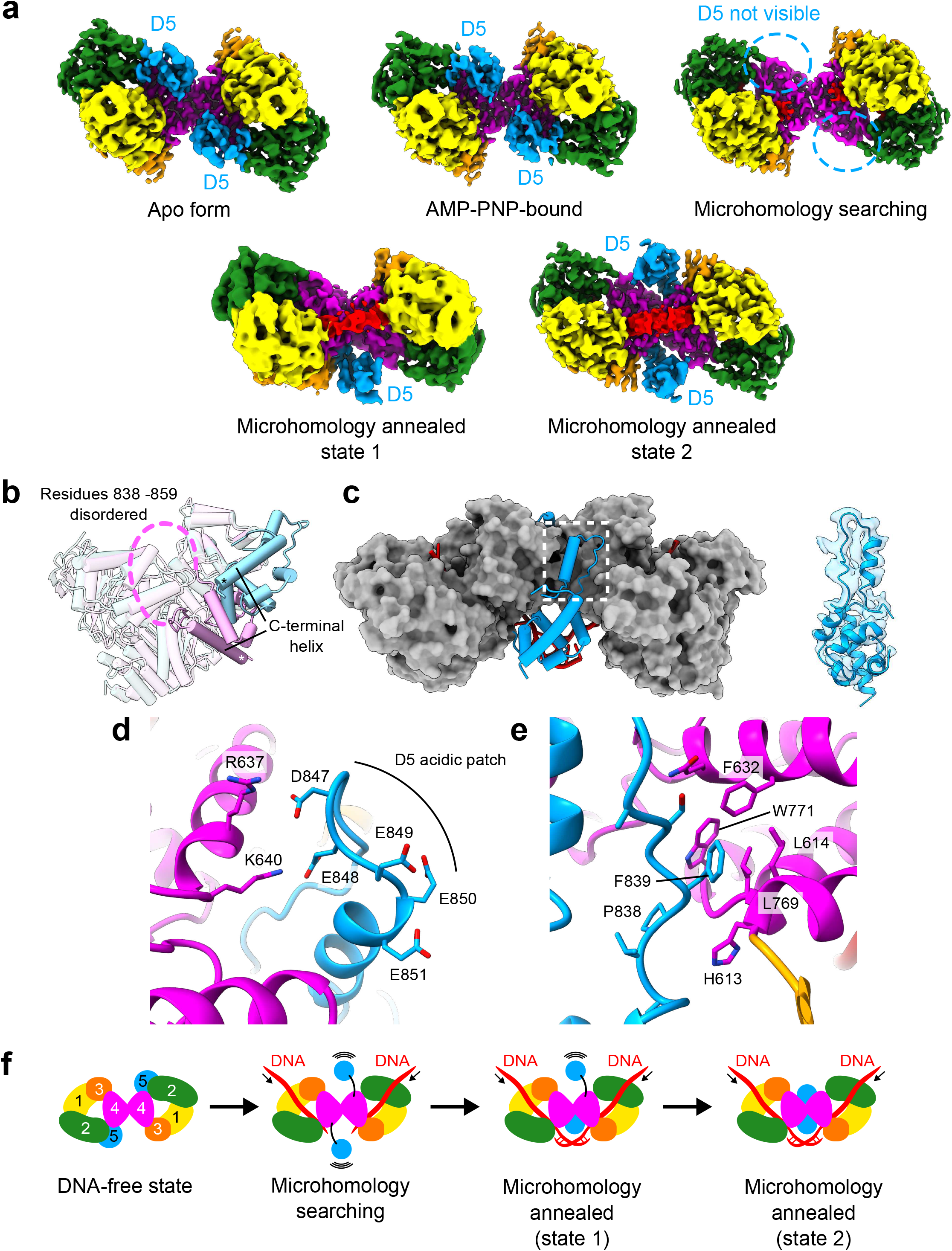
Structural variability of the mobile domain D5. (**a**) Cryo-EM density maps of Polθ-hel in the apo form, AMP-PNP-bound form, MH search state, and MH annealed states 1 and 2. The subdomain D5 (highlighted in blue) is bound to the helicase ring at the D2-D4 interface in the apo and AMP-PNP-bound forms but becomes invisible in the MH search state. In the MH annealed states, the D5 reemerges at the dimer interface in either one protomer (state 1) or both protomers (state 2). (**b**) Superimposition of a protomer from the apo form with the MH annealed state, highlighting the displacement of D5. The C-terminal end of the C-terminal helix is marked with an asterisk. (**c**) Positioning of the relocated D5 in the dimer context, depicted in a cylinder model (blue) against the rest of the Polθ-hel dimer shown in the surface representation (grey). An overlay of the cryoEM density over the atomic model of the relocated D5 is shown on the right. (**d**) Structure around the acidic patch of the U-shape structure in D5. Basic residues R637 and K640 of D4 in the other protomer electrostatically engage with the acidic patch. (**e**) Structure around the F839 of the U-shape structure in D5. The F839 is bound at the hydrophobic cavity formed by L614, F632, L769, and W771 of D4 in the other protomer. (**f**) Proposed model of the mobile D5 during the MMEJ.

Interestingly, the relocated D5 is observed in only one protomer in the complex with 11-nt poly(T) DNA while the D5 in the other protomer remains invisible (termed state 1). In the complex with 9-nt poly(T) DNA, the relocated D5 is visible in both protomers. (termed state 2). These variations indicate different microhomology annealing states. Furthermore, the previously disordered residues 839-858 of D5 became fully visible, forming a U-shaped helix-turn-loop motif, along the shallow groove at the D4-D4 interface (Fig. 2c). At the apex of the U-turn, a negatively charged patch spanning the residues -^847^DEEEE^851^- is positioned next to a positively charged surface of the D4 of the other protomer, formed by R637 and K640, effectively anchoring the U-shape motif (Fig. 2d). Additionally, F839 at the beginning of the U-shape is trapped by a solvent-exposed hydrophobic cavity formed by residues L614, F632, L769, and W771 in the D4 of the other protomer (Fig. 2e). The U-shape motif solidifies the groove at the dimer interface, doubling the dimer interface area from 951 Å^2^ in the apo form to 1956 Å^2^ in the MH annealed state 2.

The aforementioned structural observation demonstrates that Polθ-hel D5 is a mobile domain, dissociating from the helicase ring upon DNA binding to create the necessary space for the MH search at the dimer cleft. Indeed, the MH DNA would clash with the D5 at the original location in the apo form. When binding to substrate DNA, the D5 appears to relocalize to the dimer interface in a step-wise manner during the microhomology annealing (Fig. 2f). This repositioning of D5 stabilizes the dimer configuration, which may be essential for maintaining the MH annealed DNA to support the downstream steps of MMEJ.

### DNA capture by Polθ-hel channel

The 3′-overhang ssDNA is threaded through the entire central channel of the Polθ-hel ring (Fig. 3a). A continuous DNA density was observed across the channel, extending from the wide entrance formed by the subdomains D2, D3, and D4 to the narrow exit formed between the D1 and D4 (Fig. 3b and 3c). Inside the Polθ-hel ring, a total of seven and eight nucleotides of poly(T) ssDNA strand were visible for the MH search state and MH annealed state 1, respectively. The eight nucleotides observed in the MH annealed state are termed T_1_ to T_8_ (in 5′ -> 3′ direction). Throughout the channel, the ssDNA extensively interacts with three subdomains of Polθ-hel, namely D1, D2, and D4. The 5′-end of the ssDNA is held by D2 at the entrance of the helicase channel through its interaction with the phosphate backbone, facilitated by electrostatic interactions from two basic residues K348 and K347 with the 5′-phosphate of T_1_ and T_2_, respectively.

**Fig. 3.**
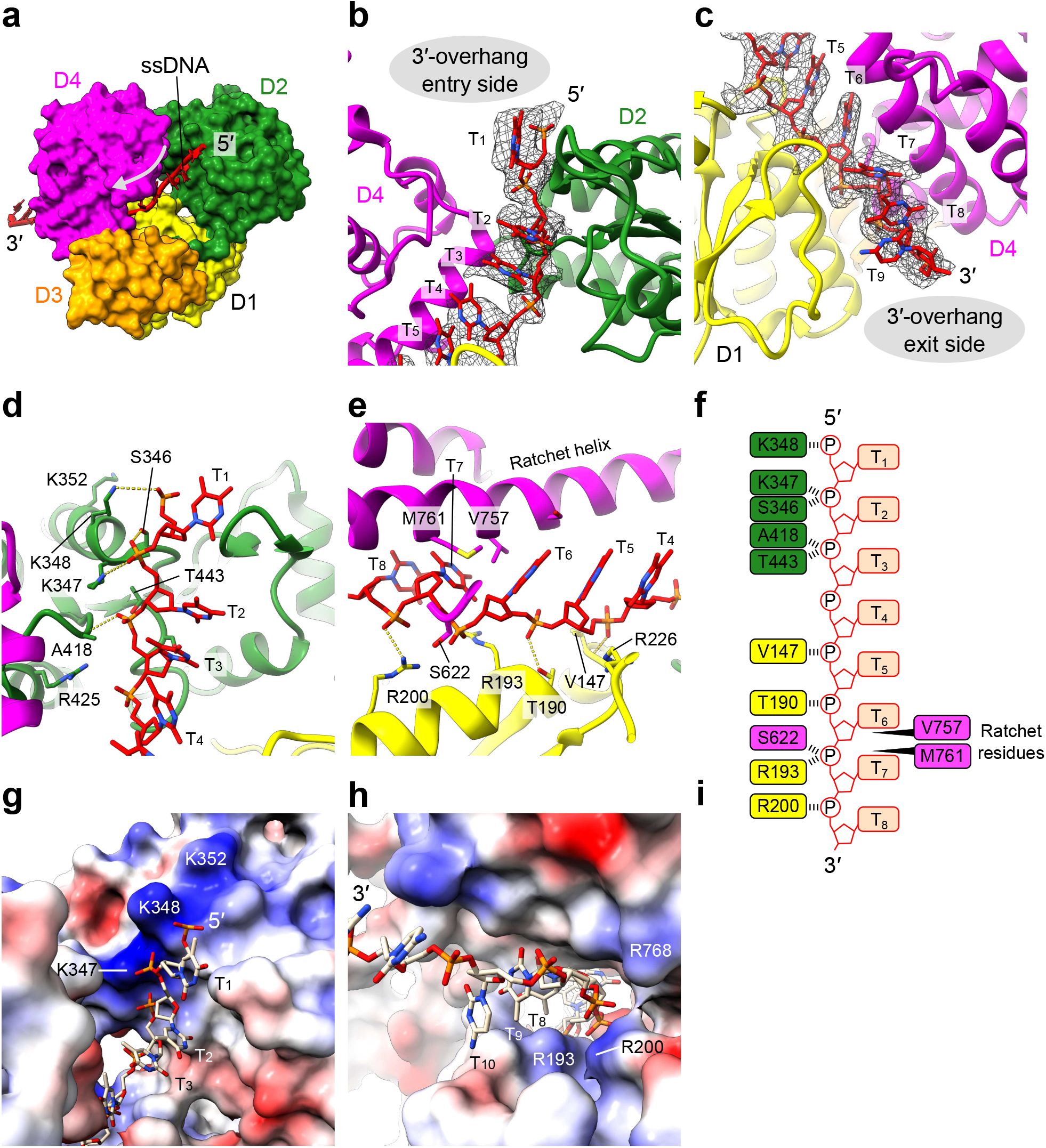
DNA-helicase channel interactions. (**a**) Overview of the 3′-overhang ssDNA (rendered as tube/slabs) threading through the helicase channel (surface model color-coded by subdomains). The direction of DNA translocation is indicated by an arrow. (**b** and **c**) Structure of ssDNA across the helicase channel, showing the atomic model of ssDNA (sticks) near the channel entrance (**b**) and exit (**c**), overlaid with the corresponding cryoEM density (mesh). (**d**) Detailed DNA-protein interactions around the channel entrance featuring the 3′-overhang DNA nucleotides T_1_-T_4_ (sticks) with surrounding amino acids (ribbons/sticks). (**e**) Detailed DNA-protein interactions around the channel exit featuring the 3′-overhang DNA nucleotides T_4_-T_8_ (sticks) with surrounding amino acids (ribbons/sticks). (**f**) Schematic of the ssDNA strand inside the Polθ-hel channel and interactions with residues from subdomains D1, D2, and D4. (**g** and **h**) Surface electrostatic potential of Polθ-hel. The positively charged patches near channel entrance (**g**) and exit (**h**) responsible for ssDNA capture. The surface area is colored according to the calculated electrostatic potential from -10.0 kT/e (red) to +10.0 kT/e (blue).

Additional stabilization is provided by the side chain hydroxyl of S346 and T443, and the main chain amine of A418, which form hydrogen bonds with the backbone phosphates of T_2_ and T_3_, thus aiding the DNA capture at the channel entrance (Fig. 3d).

The downstream ssDNA inside the channel stretches along a long α-helix in D4, which was previously termed ratchet helix (Fig. 2e), which facilitates unidirectional DNA translocation in the 3’->5’ direction as the DNA duplex is unwound by superfamily-2 helicase, acting like a ratchet^28^. The bases T_4_ to T_6_, adjacent to the ratchet helix, maintain continuous base-stacking, while two hydrophobic residues, V757 and M761 on the ratchet helix, are intercalated between the T_6_ and T_7_ bases, disrupting the base-stacking. These residues act as a wedge of the ratchet at the channel’s narrow exit, allowing unidirectional passage of the incoming DNA and preventing back-tracking. Although there is no sequence-specific interaction for the bases in this region, the phosphate backbone engages in extensive interactions with the residues in D1. The 5′-phosphates of T_5_ and T_6_ form hydrogen bonds with the main chain amine of V147 and the side chain hydroxyl of T190, respectively. The 5′-phosphates of T_7_ and T_8_ form electrostatic interactions with two arginines R193 and R200 on a short helix in D1 at the channel exit (Fig. 3e). The side chain hydroxyl of S622 additionally stabilizes the T_7_ phosphate. In total, 10 residues are involved in recognizing the backbone of the ssDNA inside the channel (Fig. 3f). The nucleotides after T_8_ are exposed to a space outside the channel, allowing the ssDNA chain to proceed to the dimer cleft for the subsequent microhomology sequence searching and annealing. The surface electrostatic potential of the Polθ-hel channel showed that both ends of the channel have positively charged surfaces for DNA capture. The K347 and K348 at the entrance form a continuous basic patch with K352 that likely contributes to the recognition of the additional DNA backbone at the branch of the fork DNA (Fig. 3g). At the exit, three arginines R193, R200, and R768 hold the ssDNA after it passes the narrowest point of the channel at the V757/M761 wedge (Fig. 3h).

### DNA binding induces dimer conformational change

Structural superimposition of the DNA-bound and apo forms of Polθ-hel dimer further revealed marked conformational rearrangements. Notably, in both the MH search state and MH annealed state, the relative positions of the two protomers within the dimer shift to an <open= form, where the two helicase rings rotate away from each other around the D4-D4 dimer interface. This movement results in the largest movements in the D2 among the Polθ-hel subdomains as it is located farthest from the dimer interface. In the MH search state, aligning D4 of one protomer (protomer 1) for superimposition shows a shift in D2 of the other protomer (protomer 2) with an r.m.s.d. of 13 Å. This shift increases to 20 Å in the MH annealed state, highlighting a global conformational shift in the dimer configuration as a result of DNA binding.

Additionally, smaller movements within the same protomer were observed, with the D2 from the same protomer rotating away from the other protomer by 3 Å in both the MH search and annealed states (Fig. 4a). These observations suggest that both inter- and intra-protomer conformational change are induced upon DNA binding. These conformational changes flatten the overall dimer architecture and create a wider space in the dimer cleft where the microhomology searching and annealing occur.

**Fig. 4.**
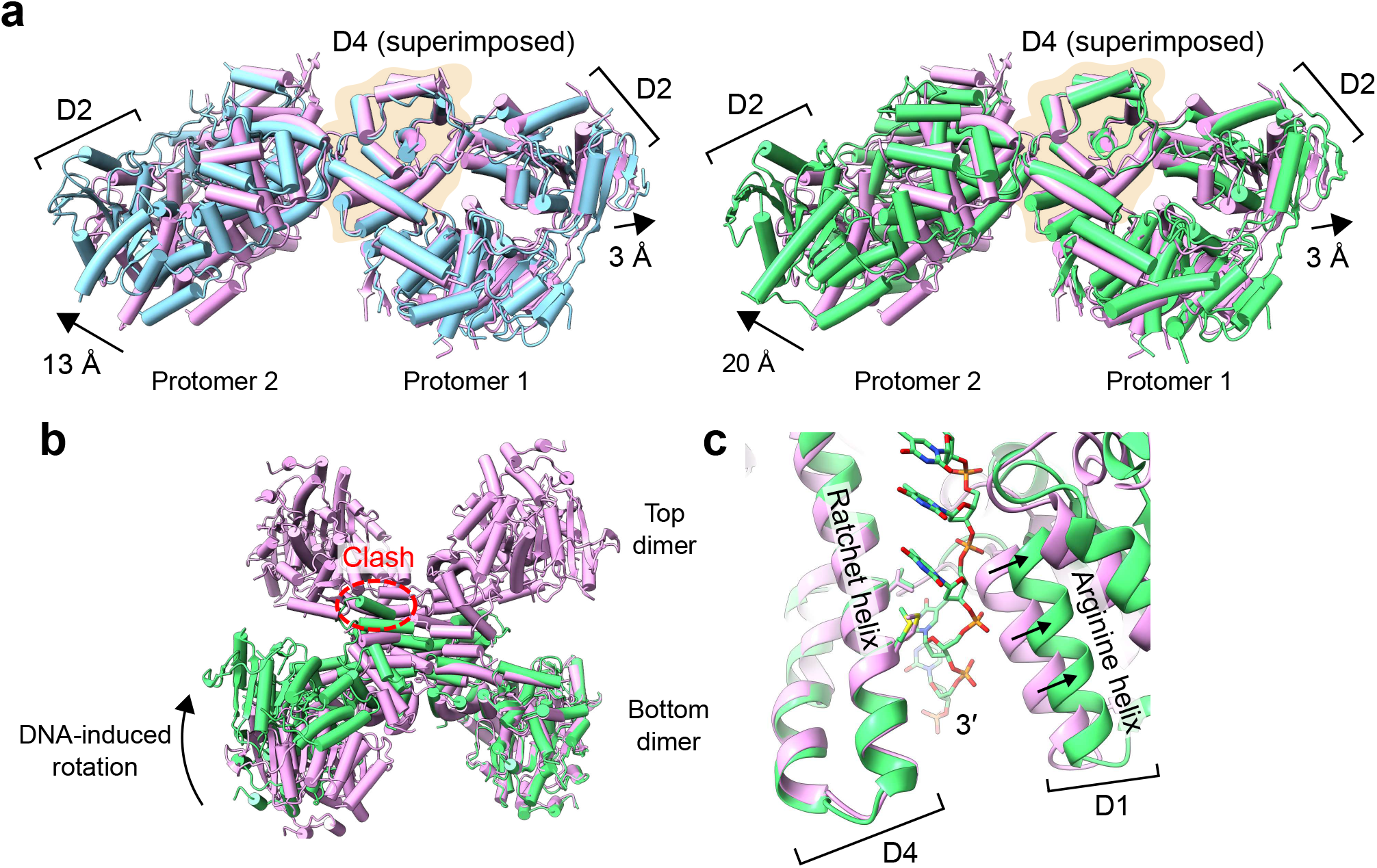
DNA-induced conformational changes in Polθ-hel dimer. (**a**) Superimposition of the Polθ-hel dimer structure in the apo form (cylinder model in light pink) and the complexes with DNA in MH search state (light blue, left) and MH annealed state (lime, right). The bound DNA and the mobile domain D5 have been omitted from the models for clarity. (**b**) Superimposition of the Polθ-hel DNA complex in MH annealed state 1 dimer form with the dimer unit of the apo form tetramer. A steric clash between the DNA-bound dimer and the other dimer unit of the apo form tetramer is highlighted. (**c**) Superimposition of the apo form (ribbon model in light pink) and the MH annealed state (lime) near the exit of the helicase channel. The movement of the arginine helix in D1 is indicated by arrows. The ratchet helix was used for alignment in the superimposition.

The superimposition of the DNA-bound dimer with one of the dimer units of the apo form tetramer structure shows a steric clash with the other dimer unit near the dimer-dimer interface, suggesting that the DNA-bound dimer conformation is incompatible with forming a tetramer as in the apo form (Fig. 4b). This is consistent with our finding in our cryoEM data sets that tetramer is only present as a DNA-free form.

As part of the intra-protomer conformational changes, the narrow exit of the DNA channel formed between the D1 and D4 widens upon DNA binding. When the ratchet helix in D4 is aligned, the helix in D1 containing two arginines (R193 and R200) that recognize backbone phosphates (termed arginine helix) shifted away from the rachet helix by 4 Å (Fig. 4c). This shit indicates a degree of flexibility in the channel exit as the ssDNA passes through it. Collectively, these results indicate that Polθ-hel undergoes both global and local conformational changes upon DNA binding, revealing that the dimer form is the functional oligomeric unit during the MMEJ process.

## Discussion

Our study elucidates the critical role of Polθ-hel in the MMEJ repair of DSBs. The cryo-EM structures of Polθ-hel, in its apo form, AMP-PNP-bound form, and DNA-bound states, elucidate the mechanism by which this helicase recognizes and processes 3′-ssDNA overhangs, facilitating microhomology search and annealing. The results reveal an unprecedented stepwise and sequential conformational change involved in the initial steps of MMEJ by Polθ-hel. The implications of these findings not only provide basic mechanistic insights but also offer specific potential drug-target sites for therapeutic intervention in HRD cancers.

Our structural analysis revealed that the active form of Polθ-hel is a dimer, mediated by head-to-head interactions of the D4 domains. Upon DNA binding, Polθ-hel undergoes notable conformational changes that transition the dimer from a “closed” to an “open” state. This shift flattens the overall dimer architecture, creating a wider cleft that accommodates the microhomology search and annealing process. The DNA-bound dimer configuration is incompatible with the tetrameric form observed in the apo state, indicating that DNA binding induces a functional dimerization that is essential for MMEJ.

A striking finding of our study is the dynamic behavior of the D5 domain which is manifested by its flexibility of location and induced refolding. In its apo form or ANP-PMP-bound form, D5 adheres to the helicase ring, but upon DNA binding, it detaches from the helicase ring and relocates to the dimer interface in a stepwise fashion, accompanied by structural refolding. This relocation of D5 is crucial for generating space for the 3′-overhang ssDNA exit to perform microhomology search near the ssDNA exits of Polθ-hel dimer interface, and the accompanied refolding is also critical for facilitating the tetramer-to-dimer transition and further solidifying the dimer interface for the MMEJ process. These overall and local conformational changes in Polθ-hel associated with DNA binding ensure the precise alignment of DNA overhangs, which is critical for the MMEJ pathway.

The detailed examination of ssDNA threading through the Polθ-hel channel reveals extensive interactions with residues across subdomains D1, D2, and D4. The ssDNA going through the entire helicase channel passage in the absence of ATP and magnesium ion suggests that the helicase ring is sufficiently flexible to allow the full traverse of its channel by the ssDNA. The interactions of Polθ-hel with the DNA are non-sequence specific with a mixture of the charge-charge interactions with the backbone and hydrophobic interactions with the bases, which stabilize the ssDNA and facilitate its unidirectional translocation. This unidirectional movement is critical for the helicase’s function, allowing it to align and anneal microhomologies efficiently.

Based on the available data, we propose a working model of Polθ-hel function in MMEJ during DSB repair (Extended Data Fig. 6). Polθ operates as a dimer mediated via Polθ-hel interactions. Consistent with this model, prior electron micrographs observed that full-length Polθ behaved mostly as monomeric and dimeric forms^10^. The Polθ-hel dimer binds to resected 3′-overhangs and dissociates replication protein A (RPA), thereby exposing the microhomology sequences necessary for end alignment and annealing.

Following this annealing, the polymerase domain of Polθ extends the annealed DNA ends sequentially and completes the repair process with the help of DNA ligases and other cofactors. The long flexible central domain linker tethers the helicase and polymerase domains and may also function to interact and coordinate with other protein cofactors necessary for MMEJ.

In summary, our findings provide crucial insights into the molecular mechanisms of Polθ-hel in DNA repair. The structural elucidation of its interactions with ssDNA and the conformational flexibility of its subdomains and its dimer/tetramer form underscore the enzyme’s structural plasticity in facilitating MMEJ in repairing DSBs. The detailed understanding of Polθ-hel’s mechanism also opens avenues for inhibitors that target specific sites of Polθ-hel to exploit synthetic lethality in HRD tumors as precision medicine for cancer therapy.

## Methods

### Protein expression and purification

The pSUMO expression vector carrying human DNA polymerase θ helicase domain (residues 1-894) was transformed into the *E. coli* strain Rosetta 2 (DE3) pLysS. The bacterial cells harboring the expression vectors were grown in LB medium at 37°C until the OD_600_ reached 0.3. The recombinant proteins were induced by 1 mM isopropyl β-D-1-thiogalactopyranoside (IPTG) at 18°C for 18 hours. The cell pellets were resuspended with a lysis buffer (25 mM Tris-HCl (pH 8.5), 500 mM NaCl, 10% glycerol, and 0.5 mM TCEP) supplemented with 2 mM PMSF and a cOmplete inhibitor tablet per 100 ml, lysed by sonication, and cellular debris was removed by centrifugation. The supernatant containing the His_6_-SUMO-Polθ-hel was loaded onto a Ni-NTA agarose column (Qiagen). The nickel column was extensively washed with a wash buffer (25 mM Tris-HCl (pH 8.5), 500 mM NaCl, 10% glycerol, 40 mM imidazole, and 0.5 mM TCEP). The His_6_-SUMO tag was cleaved with Prescission protease in one bed volume of the lysis buffer by incubating at 4°C overnight. The tag-free Polθ-hel was eluted from the resin in three bed volumes of the lysis buffer (Cytiva). The eluent was diluted with a 2.5 times larger volume of a buffer with no salt (25 mM Tris-HCl (pH 8.5), 10% glycerol, and 0.5 mM TCEP), and loaded onto the HiTrap Heparin HP affinity column (Cytiva). The proteins were eluted with a gradient NaCl of 0.2 to 2.0 M. Final purification was achieved with a Superdex 200 Increase 10/300 GL column (Cytiva) equilibrated with a buffer (25 mM Tris-HCl (pH 8.5), 500 mM NaCl, and 0.5 mM TCEP). The peak fractions were collected and concentrated for cryoEM study.

Protein purity was assessed by SDS-PAGE at each purification step.

### CryoEM sample preparation and data acquisition

Five data sets were collected in separate TEM sessions. Prior to the reconstitution of the Polθ-hel-DNA complexes, synthetic ssDNA strands (Integrated DNA Technologies) were pre-annealed to form dsDNA with 3′-overhang ssDNA. The following pairs of ssDNA were used for annealing: 9-nt poly(T) with MH 3′-overhang (chain 1: 5′-CCAACCGACCACACCCACCACCCTACCGCCTTTTTTTTTCCCGGG-3′, chain 2: 5′-GGCGGTAGGGTGGTGGGTGTGGTCGGTTGG-3′), 11-nt poly(T) with MH 3′-overhang (chain 1: 5′-CCAACCGACCACACCCACCACCCTACCGCCTTTTTTTTTTTCCCGGG-3′, chain 2: 5′-GGCGGTAGGGTGGTGGGTGTGGTCGGTTGG-3′), and 15-nt poly(T) without MH 3′-overhang (chain 1: 5′-CCAACCGACCACACCCACCACCCTACCGCCTTTTTTTTTTTTTTT-3′, chain 2: 5′-GGCGGTAGGGTGGTGGGTGTGGTCGGTTGG-3′). These pairs of ssDNA were mixed in an annealing buffer (10 mM HEPES (pH 7.5) and 50 mM NaCl), denatured at 95°C for 5 minutes, and then cooled down slowly overnight. For the data set of the complex with 9-nt poly(T) with MH, 11-nt poly(T) with MH, and 15-nt poly(T) without MH 3′-overhang DNA, the purified protein (10 μM) and pre-annealed DNA were mixed by 1:4, 1:10, and 1:30 molar ratio, respectively, in a buffer (10 mM HEPES (pH 7.5), 150 mM NaCl, and 0.5 mM TCEP) and the mixture was incubated on ice overnight. For the AMP-PNP-bound form, the purified protein (10 μM) was mixed with 2 mM AMP-PNP and the mixture was incubated on ice for 20 min. Of note, the samples for the data sets of the AMP-PNP-bound form and apo forms include DNA substrates with 3′-overhang and internal loop structures, respectively, but these samples did not yield any DNA-bound Polθ-hel structure. 3 ul aliquots of the mixture were applied to UltrAu foil R1.2/1.3 gold 300-mesh grids (Electron Microscopy Sciences). Grids were then blotted and vitrified in liquid ethane cooled by liquid nitrogen using Vitrobot Mark IV (Thermo Fisher Scientific).

CryoEM data of Polθ-hel-DNA complexes were collected in a Titan Krios G3i (Thermo Fisher Scientific) equipped with a K3 direct electron detector and post-BioQuantum GIF energy filter (Gatan) operated at 300 kV in electron counting mode. Movies were collected at a nominal magnification of 105,000× in super-resolution mode after binning by a factor of 2, resulting in an effective pixel size of 0.86 Å. A total dose of 65 e^-^/Å^2^ per movie was used with a dose rate of approximately 15 e^-^/pix/sec. 10,005, 10,331, and 7,268, movies were recorded for the complexes with 3′-overhang DNA with 9-nt poly(T) with MH, 11-nt poly(T) with MH, and 15-nt poly(T) without MH data sets, respectively, by automated data acquisition with EPU version 3.5.0.

CryoEM data of AMP-PNP-bound form and apo form of Polθ-hel were collected in a Glacios (Thermo Fisher Scientific) equipped with Falcon-4 direct electron detector operated at 200 kV in electron counting mode. Movies were collected at a nominal magnification of 150,000× and a pixel size of 0.92 Å in EER format. A total dose of 50-60 e^-^/Å^2^ per movie was used with a dose rate of 5-6 e^-^/pix/sec. 671 and 4,511 movies were recorded for AMP-PNP-bound form and apo form, respectively, by automated data acquisition with EPU.

### CryoEM data processing

The movies from five data sets were imported into cryoSPARC software package^29^ and subjected to patch motion correction and CTF estimation in cryoSPARC. Initially, reference-free manual particle picking in a small subset of data was performed to generate 2D templates for auto-picking and to assess the data quality.

For the complex with 9-nt poly(T) with MH 3′-overhang DNA, a total of 3,203,827 particles were picked initially, extracted, and down-sampled by a factor of 4, on which 2D classification was performed. 746,417 particles from 2D class averages were selected and re-extracted with full resolution. Another round of 2D classification was performed and 514,557 particles from 2D class averages were selected. 3D *ab initio* reconstruction was then performed to generate six initial volumes. The particles from the first round of 2D classification were then used in the following heterogeneous refinement with two copies of each of the six *ab initio* classes as starting volumes. A single dominant class containing 21.5% of the particles showed a feature of dimeric Polθ-hel with anisotropic density. Further *ab initio* reconstruction and heterogeneous refinement were performed with two classes to obtain more isotropic maps. A single class containing 49.2% of the particles was selected, and non-uniform refinement^30^ was performed with C2 symmetry to yield the final 3.1 Å resolution map. A single class containing 6.6% of the particles in the first heterogeneous refinement showed a feature of tetrameric Polθ-hel and it was used for non-uniform refinement with D2 symmetry to yield a 3.4 Å resolution map. The tetramer map displayed no DNA density.

For the complex with 11-nt poly(T) with MH 3′-overhang DNA, the micrographs were curated and 34% of them were discarded. A total of 1,050,266 particles were picked initially, extracted, and down-sampled by a factor of 4, on which 2D classification was performed. We noticed that free DNA particles were dominant in this data set, which interfered with the identification of protein particles. Topaz, a convolutional neural-network-based particle-picking program^31^, was trained with the clean 31,276 particles from the 2D classification. 885,890 particles were extracted by Topaz and used for the second round of 2D classification. A subset of good 2D class averages with clear secondary structure features containing 65,577 particles was used for *ab initio* reconstruction to generate four initial volumes. Two similar classes containing 64.1% of the particles with a feature of dimeric Polθ-hel were then combined to yield an *ab initio* volume. The three volumes were used as starting volumes for the following heterogeneous refinement with 105,752 particles from a broader selection of 2D classes in the second round of 2D classification. A single dominant class containing 50.7% of the particles was selected, and non-uniform refinement was performed with C1 symmetry to yield the final 3.8 Å resolution map.

For the complex with 15-nt poly(T) without MH 3′-overhang DNA, a total of 2,654,648 particles were picked initially, extracted, and down-sampled by a factor of 4, on which 2D classification was performed. A subset of 2D class averages containing 210,199 particles was re-extracted with full resolution and used for 3D *ab initio* reconstruction to generate six initial volumes. A broader selection of 2D classes containing 702,967 particles was then used in the following heterogeneous refinement with two copies of each *ab initio* class as starting volumes. A single dominant class containing 14.5% of the particles was selected, and non-uniform refinement was performed with C2 symmetry to yield the final 3.2 Å resolution map.

For the complex with AMP-PNP, a total of 501,459 particles were picked initially, extracted, and down-sampled by a factor of 4, on which 2D classification was performed. 232,215 particles from 2D classes with clear features were selected and re-extracted with full resolution. 3D *ab initio* reconstruction was then performed to generate six initial volumes. A single dominant class with a feature of dimeric Polθ-hel containing 30.3% of the particles was selected, and non-uniform refinement was performed with C2 symmetry to yield the final 3.5 Å resolution map.

For the apo form dimer and tetramer, a total of 3,882,670 particles were picked initially, extracted, and down-sampled by a factor of 4, on which 2D classification was performed. 1,965,226 particles from 2D classes were selected and re-extracted with full resolution. 3D *ab initio* reconstruction was then performed to generate six initial volumes. Heterogeneous refinement was then performed with two copies of each of the six *ab initio* classes as starting volumes. A single class with a feature of dimeric Polθ-hel containing 14.7% of the particles was selected, and non-uniform refinement was performed with C2 symmetry to yield the final 3.6 Å resolution map of the apo form dimer. Another class with a feature of tetrameric Polθ-hel containing 14.6% of the particles was selected for further *ab initio* reconstruction and heterogeneous refinement with two classes. A dominant class containing 73.0% of the particles was selected and non-uniform refinement was performed with D2 symmetry to yield the final 3.5 Å resolution map of the apo form tetramer. All resolution evaluation was performed based on the gold-standard criterion of FSC coefficient at the 0.143^32^.

### Atomic model building

An atomic model derived from crystal structures of Polθ-hel (PDB ID: 5AGA) was docked into the cryo-EM maps of Polθ-hel apo form dimer and Polθ-hel in complex with the 15-nt poly(T) without MH 3′-overhang DNA using UCSF Chimera^33^. The apo form tetramer and AMP-PNP-bound form were built based on the apo form dimer model. The complex with the 9-nt poly(T) with MH 3′-overhang DNA was built based on the complex with the 15-nt poly(T) without MH 3′-overhang DNA model. The models were initially adjusted manually to match the density map using COOT^34^ and refined with the phenix.real_space_refine module in Phenix with secondary structure restraints and geometry restraints^35-37^. The residues 839-858 in D5 of the complex with the 9-nt poly(T) with MH 3′-overhang DNA were built de novo. For the Polθ-hel-DNA complexes, well-defined nucleotide densities inside the channel facilitated the DNA model building process. DNA densities outside the Polθ-hel channel including the density for the annealed microhomology are less well-defined. A standard B-form DNA duplex of six base pairs was placed into the low-resolution density connecting the channel exit of the two protomers in the complex with the 9-nt poly(T) with MH 3′-overhang DNA and then connected to the ssDNA chains threading through the channels. The final atomic models were validated using the comprehensive cryo-EM validation tool implemented in Phenix (Table 1)^38^. All structural figures were generated with UCSF ChimeraX^39^.

### DNA binding assay by native PAGE and fluorescence anisotropy

For DNA binding assay by native gel shift, the annealed DNA (Fig. 1b) at 20 nM was titrated by Polθ-hel in 20 μl reaction volume containing 10 mM HEPES (pH 7.5), 150 mM NaCl, 0.5 mM TCEP, and 10% glycerol. Reaction mixtures were incubated on ice for 20 min and analyzed by 8% acrylamide native PAGE. The electrophoresis was performed at a constant 150 V for 75 min using 0.5x TBE. SYBR™ Gold Nucleic Acid Gel Stain (Thermo Fisher) was used for staining the gel. The gel images were visualized using an Amersham™ Typhoon™ Biomolecular Imager (GE Healthcare). For fluorescence anisotropy, binding reactions were performed in 20 mM Tris**–**HCl (pH 7.5), 1 mM DTT, 5 mM MgCl_2_, 5 % glycerol, 0.1 mg/ml BSA, 30 mM NaCl for at least 20 min at room temperature. Reactions contained 10 nM FAM-conjugated ssDNAs: FAM-9T+MH (oligos FAM-9T+MH/DNA-c), FAM-11T+MH (oligos FAM-11T+MH/DNA-c), and FAM-15T no MH (oligos FAM-15T no MH/DNA-c), and the indicated amounts of the Polθ-hel enzyme. A ClariostarPLUS plate reader (BMG Labtech) was used to measure fluorescence anisotropy. All experiments were performed in triplicate and plotted with ± s.d. GraphPadPrism 10 was used for plotting and *K*_d_ calculations. Sequences of the oligos used to create DNA substrates for the assay are as follows (DNA-c is used to anneal each of the FAM-labeled oligos): DNA-c: 5′-GGCGGTAGGGTGGTGGGTGTGGTCGGTTGG; FAM-9T+MH: FAM-5′-CCAACCGACCACACCCACCACCCTACCGCCTTTTTTTTTCCCGGG; FAM-11T+MH: FAM-5′-CCAACCGACCACACCCACCACCCTACCGCCTTTTTTTTTTTCCCGGG; FAM-15T no MH: FAM-5′-CCAACCGACCACACCCACCACCCTACCGCCTTTTTTTTTTTTTTTTT.

## Acknowledgements

This work was supported by National Institutes of Health grants R01AI150524 to X.S.C. and R35GM152198 to R.T.P.. F.I. was a former fellowship awardee from Nakajima Foundation and Z.L. is a recipient of University of Southern California (USC) dean’s fellowship. Electron microscopy data were collected at the Core Center of Excellence in Nano Imaging (CNI) at USC. Cryo-EM data was computed at Center for Advanced Research Computing (CARC) at USC. We thank Tomek Osinski for assisting with computing work at CARC.

## Data availability

The atomic models have been deposited in the PDB with accession codes 9C5Q (Polθ-hel-DNA MH annealed state 2 dimer), 8W0A (Polθ-hel-DNA MH search state dimer), 9ASJ (Polθ-hel-AMP-PNP dimer), 9ASK (Polθ-hel apo form dimer), and 9ASL (Polθ-hel apo form tetramer). The cryo-EM density maps have been deposited in the EMDB with accession codes EMD-45217 (Polθ-hel-DNA MH annealed state 1 dimer), EMD-45218 (Polθ-hel-DNA MH annealed state 2 dimer), EMD-43706 (Polθ-hel-DNA MH search dimer), EMD-43816 (Polθ-hel-AMP-PNP dimer), EMD-43817 (Polθ-hel apo form dimer), and EMD-43818 (Polθ-hel apo form tetramer).

## Author contributions

X.S.C., R.T.P., Z.L., and F.I. conceived the project. X.S.C. supervised the project. F.I. and Z.L. purified Polθ-hel protein and performed DNA binding and cryo-EM structural studies. H.A.K. assisted with cryo-EM data collection.

L.M. performed fluorescence anisotropy assay. F.I. and Z.L. wrote the initial draft and all the authors contributed to the final version of the manuscript.

