## Extended Data for "Structural Basis for Polθ-Helicase DNA Binding and Microhomology-Mediated End-Joining"

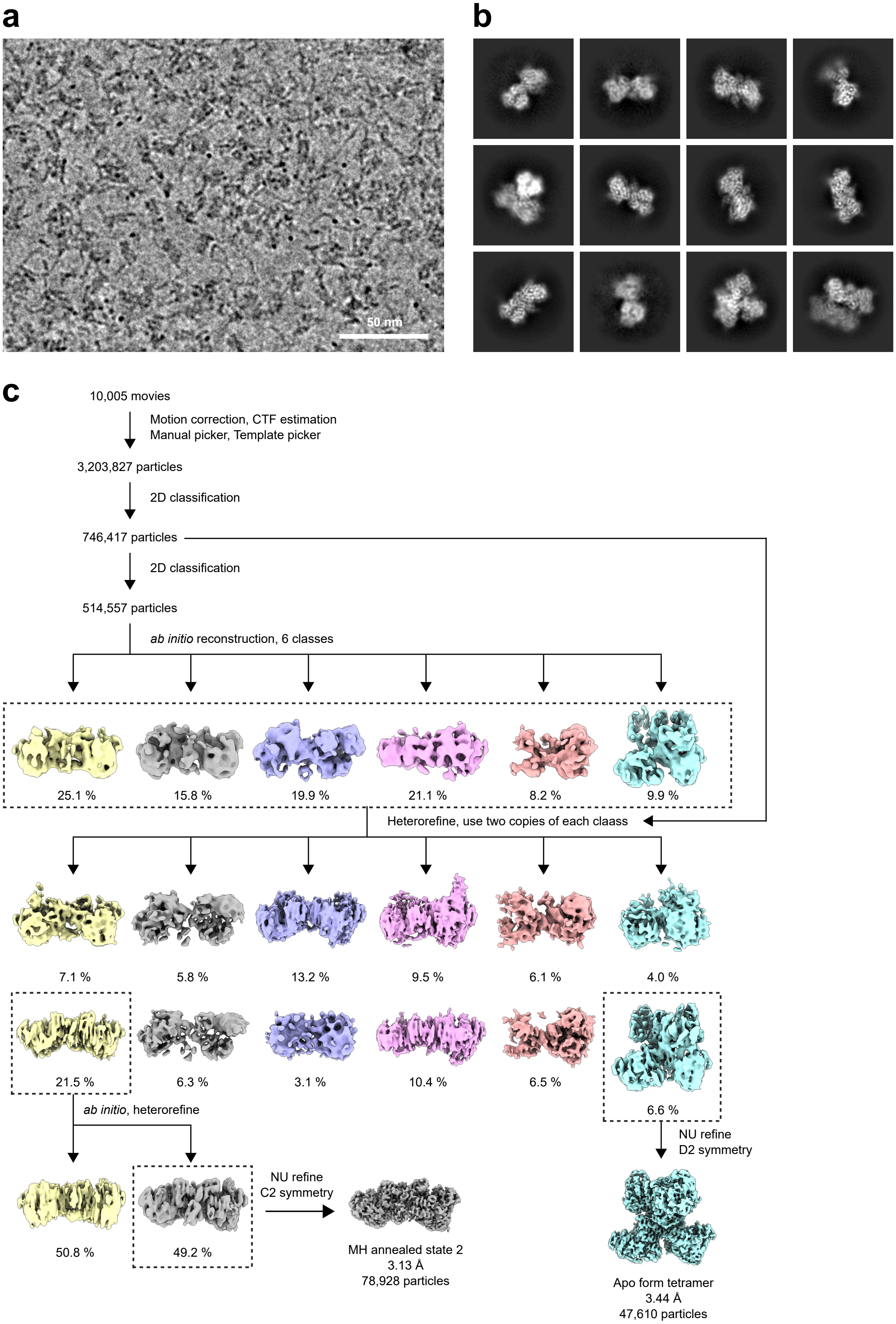

a
50 nm
b
c
10,005 movies
Motion correction, CTF estimation
Manual picker, Template picker
3,203,827 particles
2D classification
746,417 particles
2D classification
514,557 particles
*ab initio* reconstruction, 6 classes
25.1 %
15.8 %
19.9 %
21.1 %
8.2 %
9.9 %
Heterorefine, use two copies of each claass
7.1 %
5.8 %
13.2 %
9.5 %
6.1 %
4.0 %
21.5 %
6.3 %
3.1 %
10.4 %
6.5 %
6.6 %
*ab initio*, heterorefine
NU refine
D2 symmetry
50.8 %
49.2 %
NU refine
C2 symmetry
MH annealed state 2
3.13 Å
78,928 particles
Apo form tetramer
3.44 Å
47,610 particles

**Extended Data Fig. 1. Workflow and intermediate results of electron microscopy imaging and 3D reconstruction of Polθ-hel-DNA complex in MH annealed state 2.** (**a**) Representative cryo-EM raw image. (**b**) Representative 2D class averages. (**c**) Cryo-EM image processing workflow.

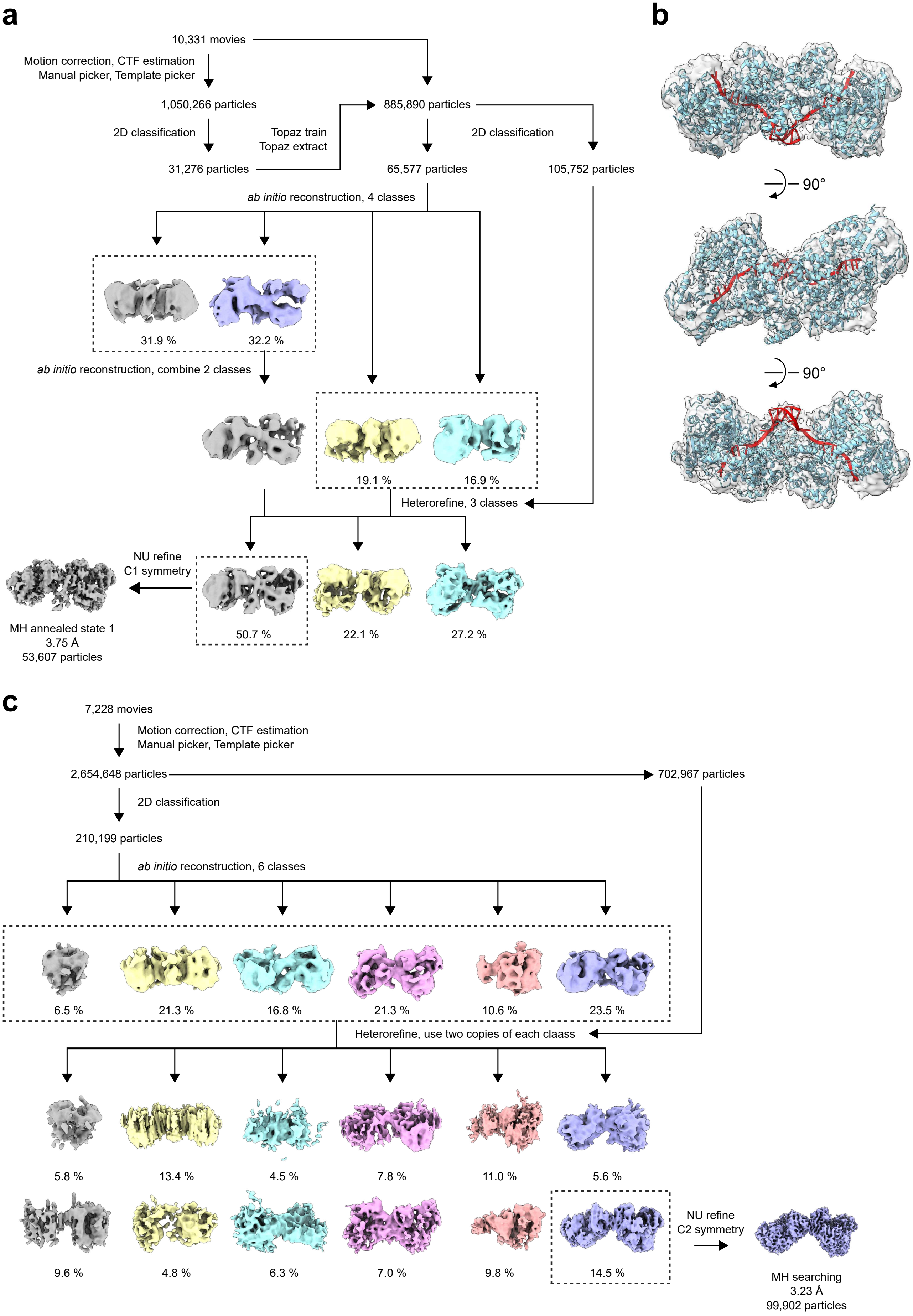
a
10,331 movies
Motion correction, CTF estimation
Manual picker, Template picker
1,050,266 particles
2D classification
31,276 particles
Topaz train
Topaz extract
885,890 particles
2D classification
65,577 particles
105,752 particles
ab initio reconstruction, 4 classes
31.9 %
32.2 %
ab initio reconstruction, combine 2 classes
19.1 %
16.9 %
Heterorefine, 3 classes
NU refine
C1 symmetry
50.7 %
22.1 %
27.2 %
MH annealed state 1
3.75 Å
53,607 particles
b
90°
90°
c
7,228 movies
Motion correction, CTF estimation
Manual picker, Template picker
2,654,648 particles
702,967 particles
2D classification
210,199 particles
ab initio reconstruction, 6 classes
6.5 %
21.3 %
16.8 %
21.3 %
10.6 %
23.5 %
Heterorefine, use two copies of each claass
5.8 %
13.4 %
4.5 %
7.8 %
11.0 %
5.6 %
9.6 %
4.8 %
6.3 %
7.0 %
9.8 %
14.5 %
NU refine
C2 symmetry
MH searching
3.23 Å
99,902 particles

**Extended Data Fig. 2. Workflow and intermediate results of electron microscopy imaging and 3D reconstruction of Polθ-hel-DNA complex in MH annealed state 1 and MH search state.** (**a**) Cryo-EM image processing workflow of the Polθ-hel-DNA complex in MH annealed state 1. (**b**) Global cryo-EM density of the Polθ-hel-DNA complex in MH annealed state 1 dimer superimposed with the atomic model derived from MH annealed state 2. The subdomain D5 in one protomer has been truncated from the model. (**c**) Cryo-EM image processing workflow of the Polθ-hel-DNA complex in MH search state.

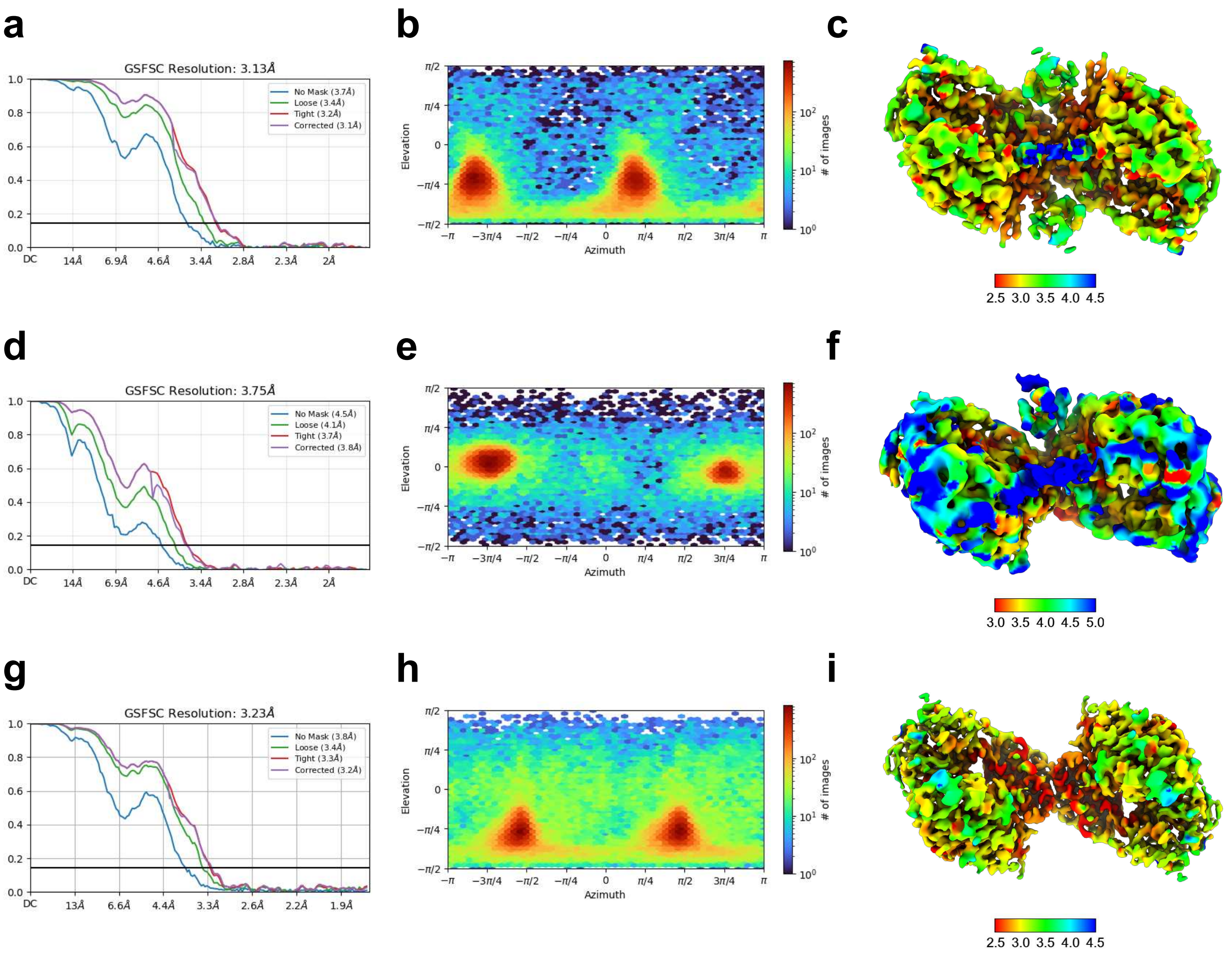


**Extended Data Fig. 3. Global resolution, particle angular distribution, and local resolution distribution of Polθ-hel-DNA complexes.** Global resolution estimation (**a**), angular distribution plot of the particles (**b**), and local resolution evaluation (**c**) of the Polθ-hel-DNA complex in MH annealed state 2. Global resolution estimation (**d**), angular distribution plot of the particles (**e**), and local resolution evaluation (**f**) of the Polθ-hel-DNA complex in MH annealed state 1. Global resolution estimation (**g**), angular distribution plot of the particles (**h**), and local resolution evaluation (**i**) of the Polθ-hel-DNA complex in MH search state. Resolution estimation is based on the gold standard Fourier shell correlation (FSC) coefficient of 0.143 criteria.

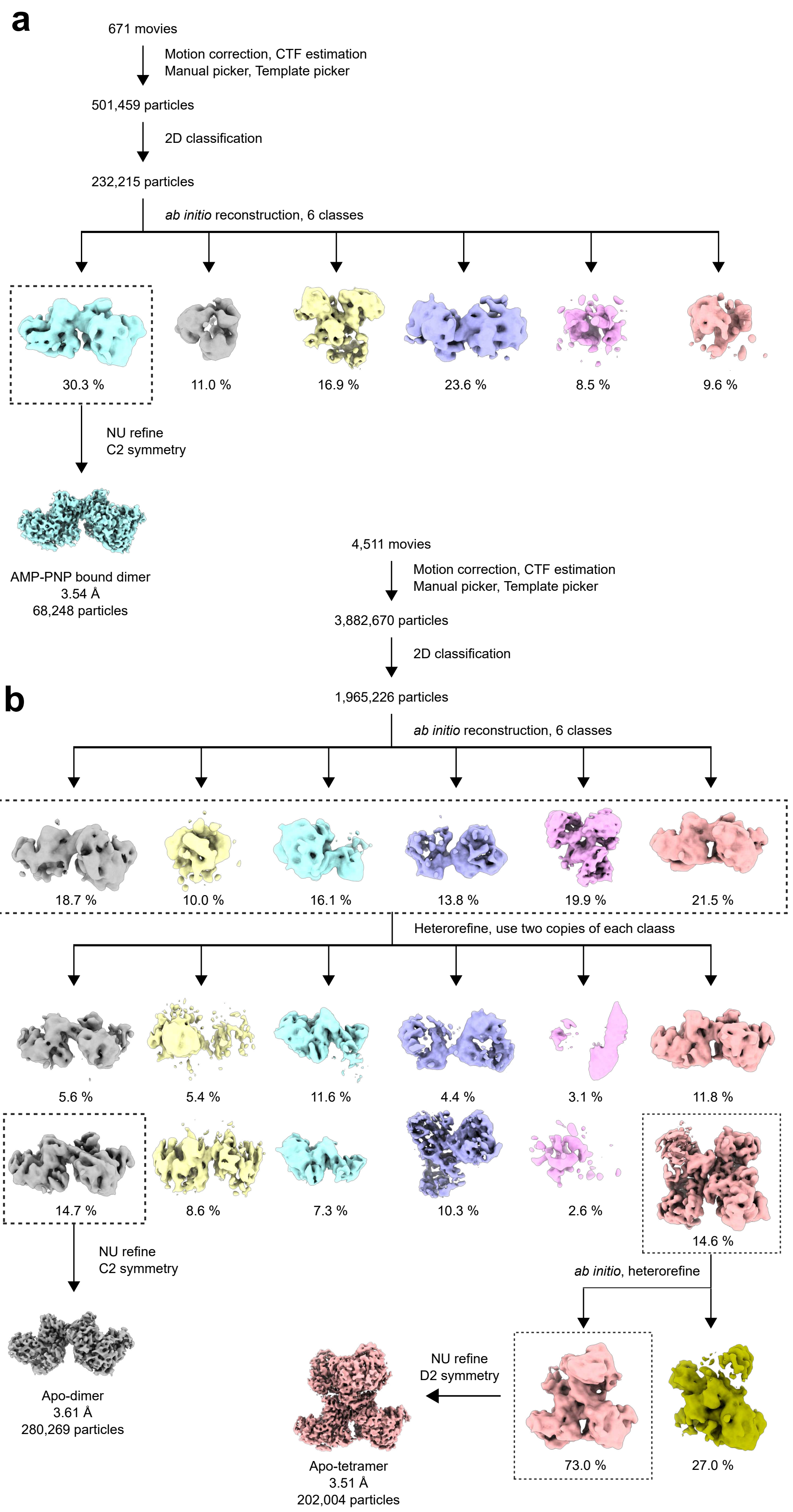
a
671 movies
Motion correction, CTF estimation
Manual picker, Template picker
501,459 particles
2D classification
232,215 particles
ab initio reconstruction, 6 classes
30.3 %
11.0 %
16.9 %
23.6 %
8.5 %
9.6 %
NU refine
C2 symmetry
AMP-PNP bound dimer
3.54 Å
68,248 particles
4,511 movies
Motion correction, CTF estimation
Manual picker, Template picker
3,882,670 particles
2D classification
1,965,226 particles
b
ab initio reconstruction, 6 classes
18.7 %
10.0 %
16.1 %
13.8 %
19.9 %
21.5 %
Heterorefine, use two copies of each claass
5.6 %
5.4 %
11.6 %
4.4 %
3.1 %
11.8 %
14.7 %
8.6 %
7.3 %
10.3 %
2.6 %
14.6 %
NU refine
C2 symmetry
ab initio, heterorefine
Apo-dimer
3.61 Å
280,269 particles
NU refine
D2 symmetry
73.0 %
27.0 %
Apo-tetramer
3.51 Å
202,004 particles

**Extended Data Fig. 4. Workflow and intermediate results of electron microscopy imaging and 3D reconstruction of Polθ-hel in complex with AMP-PNP and apo forms.** (**a**) Cryo-EM image processing workflow of the Polθ-hel-AMP-PNP complex. (**b**) Cryo-EM image processing workflow of the Polθ-hel apo form dimer and tetramer.

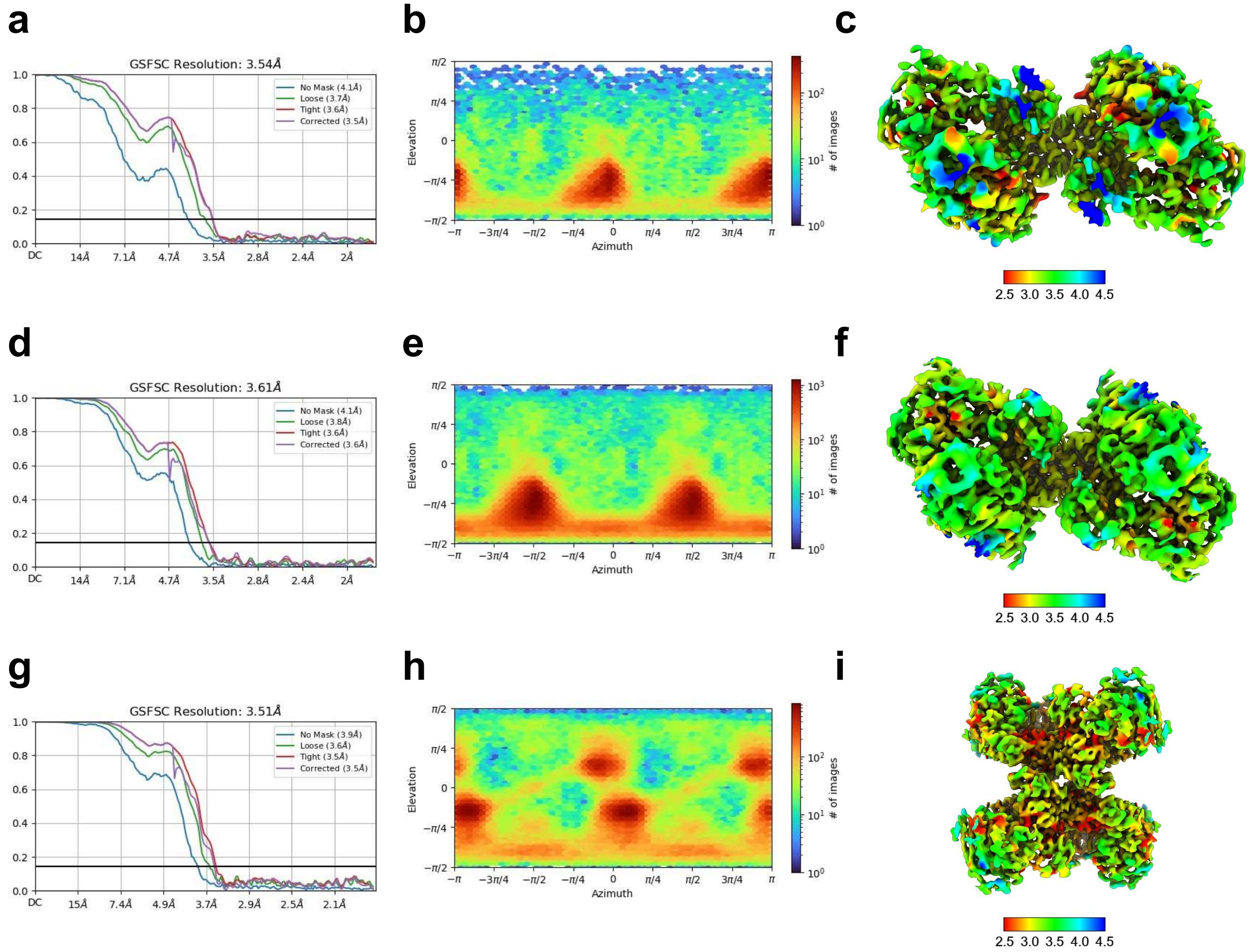


**Extended Data Fig. 5. Global resolution, particle angular distribution, and local resolution distribution of Polθ-hel in complex with AMP-PNP and apo forms.** Global resolution estimation (**a**), angular distribution plot of the particles (**b**), and local resolution evaluation (**c**) of the Polθ-hel-DNA in complex with AMP-PNP. Global resolution estimation (**d**), angular distribution plot of the particles (**e**), and local resolution evaluation (**f**) of the Polθ-hel apo form dimer. Global resolution estimation (**g**), angular distribution plot of the particles (**h**), and local resolution evaluation (**i**) of the Polθ-hel apo form tetramer. Resolution estimation is based on the gold standard Fourier shell correlation (FSC) coefficient of 0.143 criteria.

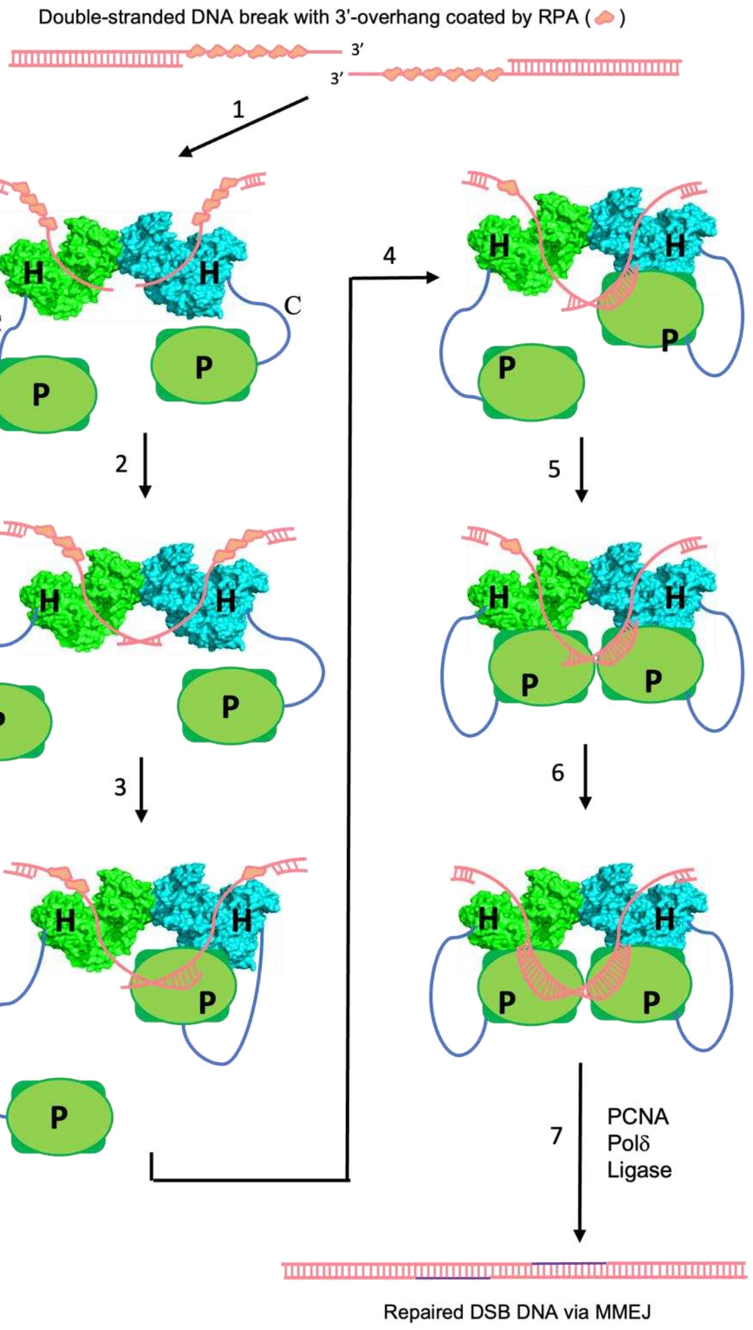


**Extended Data Fig. 6. A model of Polθ-hel roles at the initial steps in MMEJ.** The two ends of double-stranded DNA breaks (DSB) are shown at the top, with RPA binding to the resected 3′-overhang ssDNA. The Polθ forms a dimer via Polθ-hel (each protomer labeled as "H" and colored in green and cyan), with Polθ-Pol (labeled as "P") tethered to the helicase dimer through the flexible Polθ-Ctr (labeled as "C"). Step 1: Each protomer of the Polθ-hel dimer binds to one of the two resected 3′-ssDNA overhangs, bringing the 3′-ssDNA end close to each other at the dimer cleft, allowing the two 3′ end to search for microhomology sequence for base-paring. Polθ-hel binding can also displace the RPA on the ssDNA. Step 2: Polθ-hel samples the ssDNA sequence homology by translocating along the ssDNA and anneals the ssDNA ends at a location with 2-6 bp microhomology. Steps 3, 4: One Polθ-pol could then bind to the short annealed microhomology dsDNA to extend the primer. Step 5, 6: When the first Polθ-Pol extends the primer to a certain length, the second Polθ-pol can bind to extend the primer in the opposite direction to complete the polymerization. Step 7: In addition to Polθ, Polδ, PCNA, DNA ligase, and possibly other protein factors are involved in the eventual sealing of the DBS to complete the MMEJ repair.
